# The glucocorticoid response element near the sphingosine-1-phopshate receptor 3 gene mitigates inflammatory processes and maladaptive behavior in females and stressed males

**DOI:** 10.1101/2024.01.09.574910

**Authors:** Brian F. Corbett, Jay Arner, Sandra Luz, Valerie Estela-Pro, Jason Yan, Brittany Osborne, Leonel Joannas, Jorge Henao-Mejia, Jose Castro-Vildosola, Tamara Hala, Craig Bassing, Adele Harman, Deanne Taylor, Seema Bhatnagar

## Abstract

It is well established that glucocorticoid receptors (GRs) bind DNA, regulate gene expression, reduce inflammatory processes, and modulate behavior. However, the precise loci bound by GRs that are necessary for these effects are not fully understood. Here, we deleted the GR binding site near the sphingosine-1-phospate receptor 3 gene using a CRISPR/Cas9 approach (S1PR3^GR-/GR-^ rats). Defeated S1PR3^GR-/GR-^ males displayed increased inflammatory markers and social anxiety-like behavior. Similar effects were observed in non-stressed females, indicating a greater dependence for GR-induced S1PR3 in females. Coherent neural activity between the locus coeruleus (LC) and medial prefrontal cortex (mPFC) was increased in S1PR3^GR-/GR-^ males following 7 defeats. Chemogenetically inhibiting mPFC-projecting LC neurons during defeat increased subsequent social interaction in wild-type and S1PR3^GR-/GR-^ males. Together, these findings demonstrate that GR-induced S1PR3 promotes resilience by mitigating stress-induced inflammatory processes and LC-mPFC coherence.

**One Sentence Summary:** Glucocorticoid receptor-induced expression of sphingosine-1- phosphate receptor 3 reduces social anxiety-like behavior by mitigating stress-induced inflammatory processes and coherent neural activity between the locus coeruleus and medial prefrontal cortex.

## Introduction

Stress contributes to the onset and development of mood-related disorders like generalized anxiety, major depressive disorder (MDD), and post-traumatic stress disorder^1–3^, which share comorbidity with immune disorders^4–7^. Stress activates the hypothalamic-pituitary-adrenal (HPA) axis, resulting in increased levels of circulating glucocorticoids^8^. Glucocorticoids bind to glucocorticoid receptors (GRs) in peripheral tissues and the brain^9,10^. Once bound to their ligand, GRs translocate to the nucleus and bind to specific DNA sequences called glucocorticoid response elements (GREs). GRs then recruit histone-modifying enzymes to increase or decrease the expression of genes, thus regulating a wide range of biological processes^11^, including reducing inflammatory processes^12^. Inflammatory processes are increased in humans with stress-related disorders^13,14^, which may be at least partially due to glucocorticoid insensitivity caused by reduced GR expression, post-translational modifications of GRs that impair GR function^15,16^, or increased expression of FKBP5^17^, which inhibits GR function^18^.

Women, who are twice as likely to suffer from stress-related mood disorders compared to men^19^, also experience resistance to glucocorticoid-mediated reductions in inflammatory cytokines^20^. In female rodents, homeostasis of inflammatory processes may be dependent on increased levels of circulating corticosterone under baseline conditions^21,22^ as HPA axis activity is increased by estradiol^23^ and reduced by androgens^24^. Anti-inflammatory treatments reduce symptom severity in MDD^25^, especially in individuals displaying increased inflammatory biomarkers^26^. Indeed, GR knockout in the rodent forebrain causes neuroendocrine and behavioral phenotypes similar to those of depressed humans^27^. Mild GR agonism reduces PTSD symptom severity^28–30^, presumably by normalizing GR signaling as excess negative feedback of the HPA axis contributes to reduced baseline cortisol levels in individuals afflicted with PTSD^2,31^. Thus, GRs reduce inflammatory processes and mitigate stress-induced changes in behavior. However, the precise genomic binding sites of GRs that mitigate stress-induced inflammatory processes and promote stress resilience are not well understood.

We previously demonstrated that GRs are increased in the medial prefrontal cortex (mPFC) of rats that are resilient to the adverse effects of stress caused by the resident-intruder paradigm of social defeat. We showed that GR-induced sphingosine-1-phosphate receptor 3 (S1PR3, endothelial differentiation gene-3) promotes resilience by mitigating stress-induced inflammatory processes in the mPFC. S1PR3 has translational relevance as blood *S1PR3* mRNA is reduced in veterans with PTSD compared to combat-exposed controls and inversely correlates with symptom severity^32^. Here, we use a CRISPR/Cas9-based approach to develop a rat model of stress vulnerability by deleting the GR binding site that is 54,910 base pairs (bp) upstream from the transcriptional start site for the *S1pr3* gene (S1PR3^GR-/GR-^ rats). We demonstrate that compared to defeated wild-type littermates, male S1PR3^GR-/GR-^ rats display reduced S1PR3 in the mPFC, increased inflammatory markers, and social anxiety-like behavior. Similar findings are found in non-defeated females, suggesting more severe effects caused by the loss of GR-induced S1PR3. This may be because, compared to males, baseline plasma corticosterone concentrations are 5-10 times higher in females ^21,23^. We also demonstrate that coherent neural activity between the locus coeruleus (LC) and mPFC, which is increased by social defeat in wild-type males^33^, is higher in S1PR3^GR-/GR-^ rats. We show that chemogenetic inhibition of mPFC-projecting LC neurons increases social interaction in defeated wild-type and S1PR3^GR-/GR-^ males. Together, these findings show that deletion of the GRE near S1PR3 plays an important role in mitigating the adverse effects of stress.

### GRs increase S1PR3 expression in females and defeated males

We previously demonstrated that social defeat stress increases blood S1PR3 mRNA expression in resilient, but not vulnerable, rats. S1PR3 protein is increased in mPFC neurons of resilient rats compared to non-stressed controls and vulnerable rats. This is due, at least in part, to GRs, which are increased in the mPFC of resilient rats, as GR knockdown caused reduced S1PR3 protein in the mPFC of defeated rats^32^. In mice, a GR binding site has been identified near the *S1pr3* gene, but no other genes encoding S1PRs^34^. Using a BLAT alignment, we identified the homologous sequence in rats, which is 54,910 bp upstream from the transcriptional start site for S1PR3 (Fig. 1a). We used chromatin immunoprecipitation (ChIP) to demonstrate that GR binds to this sequence in the mPFC as GR binding was increased at this locus in defeated wild-type males compared to non-defeated males (Fig. 1b). No amplification was observed for S1PR3^GR-/GR-^ rats, confirming knockout of this locus. Circulating plasma corticosterone concentrations are 5-10 times higher under baseline conditions in females compared to males^21,23^. Therefore, we hypothesized that S1PR3 might be increased in wild-type, but not S1PR3^GR-/GR-^, females compared to males under baseline conditions. Male and female wild-type and S1PR3^GR-/GR-^ rats were either subjected to seven days of social defeat or served as novel cage controls. For non-defeated wild-type, but not S1PR3^GR-/GR-^, rats S1PR3 is increased in the prelimbic (PL) and infralimbic (IL) cortices of non-defeated females compared to non-defeated males. Seven days of defeat does not increase S1PR3 in the PL or IL of wild-type females, but S1PR3 is reduced in S1PR3^GR-/GR-^ rats compared to wild-type controls regardless of defeat condition. Consistent with previous findings^32^, GRs increases S1PR3 in the mPFC of males as defeat increases S1PR3 in the infralimbic cortex (IL) of male wild-type, but not S1PR3^GR-/GR-^, rats (Fig. 1c-e). We found similar effects in whole blood, as in the absence of defeat, S1PR3 mRNA was increased in wild-type, but not S1PR3^GR-/GR-^, females compared to males (Fig. 1f). In defeated males, S1PR3 mRNA was reduced in S1PR3^GR-/GR-^ rats compared to defeated wild-type littermates (Fig. 1g). Together, these findings show that corticosterone-activated GRs increase S1PR3 expression in mPFC neurons and blood.

**Figure 1.**
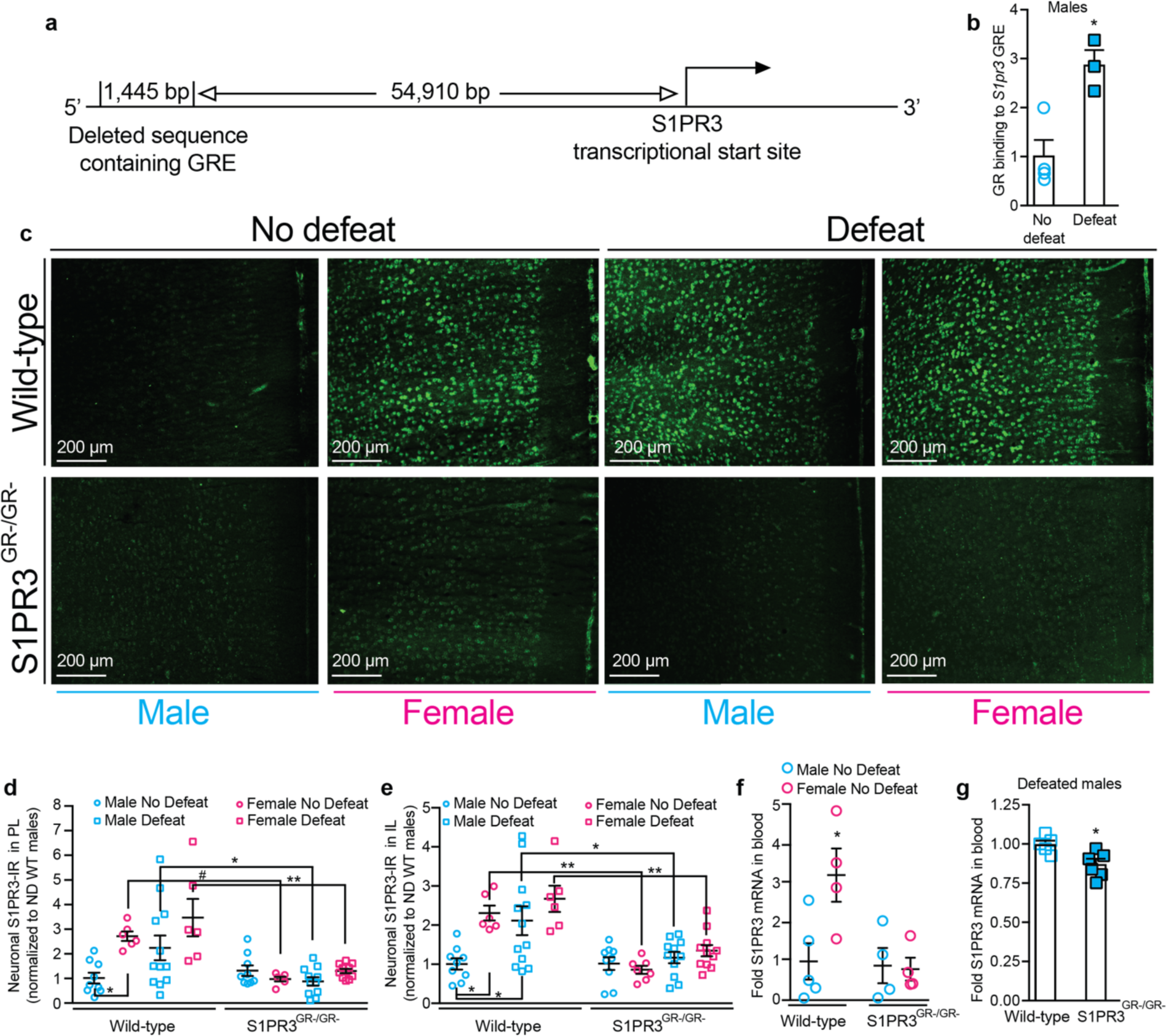
GR-induced S1PR3 is impaired in S1PR3^GR-/GR-^ rats. (**a**) Graphic model of deleted GRE in S1PR3^GR-/GR-^ rats. (**b**) GR binding to the GRE upstream of the *S1pr3* gene in non-defeated (n=3) and defeated (n=4) male wild-type rats. (**c**) Images and quantification of S1PR3 in the (**d**) PL and (**e**) IL of non-defeated wild-type males (n=9), non-defeated wild-type females (n=6), defeated wild-type males (n=12), defeated wild-type females (n=6), non-defeated S1PR3^GR-/GR-^ males (n=9), non-defeated S1PR3^GR-/GR-^ females (n=7), defeated S1PR3^GR-/GR-^ males (n=11), and defeated S1PR3^GR-/GR-^ females (n=11). (**f**) S1PR3 mRNA in whole blood taken from non-defeated wild-type males (n=5), wild-type females (n=4), S1PR3^GR-/GR-^ males (n=4), and S1PR3^GR-/GR-^ females (n=4). (**g**) S1PR3 mRNA in whole blood collected from defeated wild-type males (n=5) and defeated S1PR3^GR-/GR-^ males (n=6). Lines represent means ± SEM. For (b) and (g), *p<0.05; unpaired two-tailed Student’s t-test. For (d) and (e), *p<0.05, **p<0.01, #p<0.10; Tukey’s multiple comparisons following 3-way ANOVA. For (f), *p<0.05 compared to all other groups; Tukey’s multiple comparisons following 2-way ANOVA.

### S1PR3^GR-/GR-^ rats display vulnerable-like behavior

In the resident-intruder paradigm of social defeat stress, rats that passively cope by displaying reduced defeat latencies in response to aggression from a retired male breeder exhibit social anxiety-like behavior as subsequent social interaction with a non-aggressive conspecific is reduced^35^. We previously demonstrated that S1PR3 knockdown in the mPFC reduces defeat latencies and social interaction, indicating a more vulnerable phenotype^32^. Consistent with these results, we found that defeat latency is reduced in male S1PR3^GR-/GR-^ rats compared to wild-type littermates (Fig. 2a). This effect was not observed in females (Fig. 2b), which use a modified resident-intruder paradigm in which a lactating female is used as an aggressor. Social defeat reduced social interaction in males and females, an effect that was exacerbated in S1PR3^GR-/GR-^ rats. Social interaction was reduced in S1PR3^GR-/GR-^ females compared to wild-type female litter mates even in the absence of stress (Fig. 2c). Together, these findings indicate that GR-induced S1PR3 is necessary for stress resilience in defeated males and mitigates social anxiety-like behavior during both baseline and stressed conditions in females.

**Figure 2.**
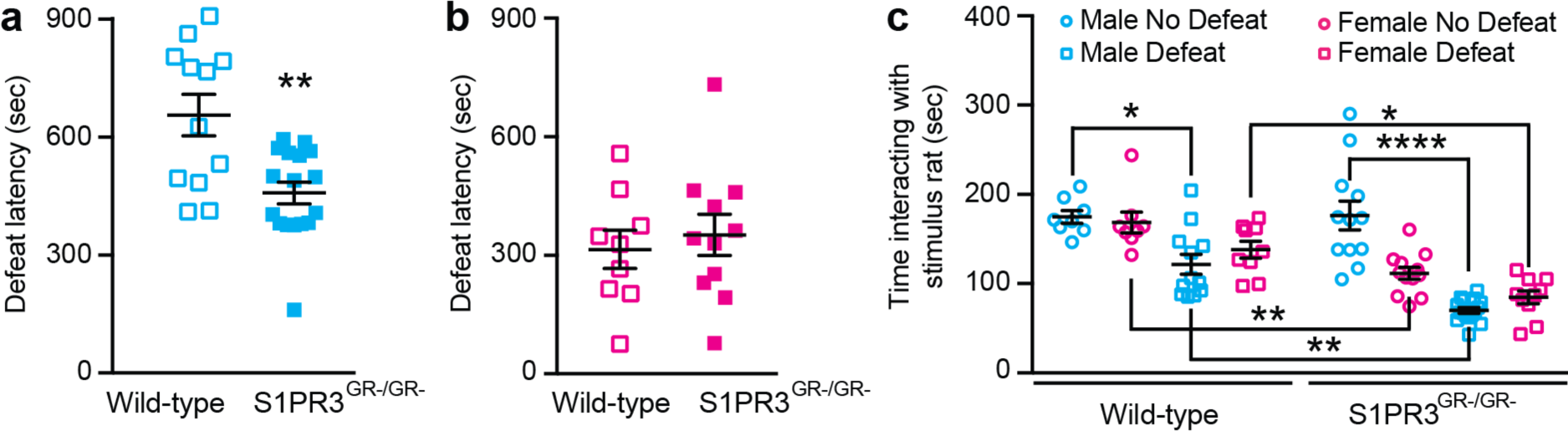
Social interaction is reduced in defeated S1PR3^GR-/GR-^ males and non-defeated S1PR3^GR-/GR-^ females. Defeat latencies are reduced in (**a**) S1PR3^GR-/GR-^ males (n=16) compared to wild-type males (n=12) but not in (**b**) S1PR3^GR-/GR-^ females (n=11) compared to wild-type females (n=9). (**c**) Time interacting with the stimulus rat in the social interaction paradigm in non-defeated wild-type males (n=8), non-defeated wild-type females (n=8), defeated wild-type males (n=12), defeated wild-type females (n=9), non-defeated S1PR3^GR-/GR-^ males (n=12), non-defeated S1PR3^GR-/GR-^ females (n=12), defeated S1PR3^GR-/GR-^ males (n=16), and defeated S1PR3^GR-/GR-^ females (n=10). Lines represent means ± SEM. For (a), **p<0.01; unpaired two-tailed Student’s t-test. For (c), *p<0.05, **p<0.01, ****p<0.0001; Tukey’s multiple comparisons following 3-way ANOVA.

### Central and peripheral inflammatory processes are increased in S1PR3^GR-/GR-^ rats

We previously demonstrated that S1PR3 mitigates defeat-induced increases in ionized calcium-binding adaptor molecule (IBA1)+ cell densities and expression of the inflammatory cytokine tumor necrosis factor alpha (TNFα) in the mPFC^32^. S1PR3 also reduces inflammatory processes in peripheral tissue^36,37^. S1PR3 may confer anti-inflammatory effects via its inhibition of nuclear factor kappa B (NFkB)^38^, the primary transcriptional regulator of inflammatory gene expression^39,40^. In the PL, IBA1+ cell densities were reduced in females (sex effect, p = 0.0227) and by defeat (defeat effect p = 0.0445) (Fig. 3a). No significant differences among groups were observed in PL. In the IL, IBA1+ cells densities were also reduced in females (sex effect, p < 0.001) and defeated S1PR3^GR-/GR-^ males displayed higher IBA1+ cell densities compared to non-defeated S1PR3^GR-/GR-^ rats, defeated wild-type males, and defeated S1PR3^GR-/GR-^ females (Fig 3b,c). This indicates that GR-induced S1PR3 mitigates stress-induced microglia densities in the IL of male rats.

**Figure 3.**
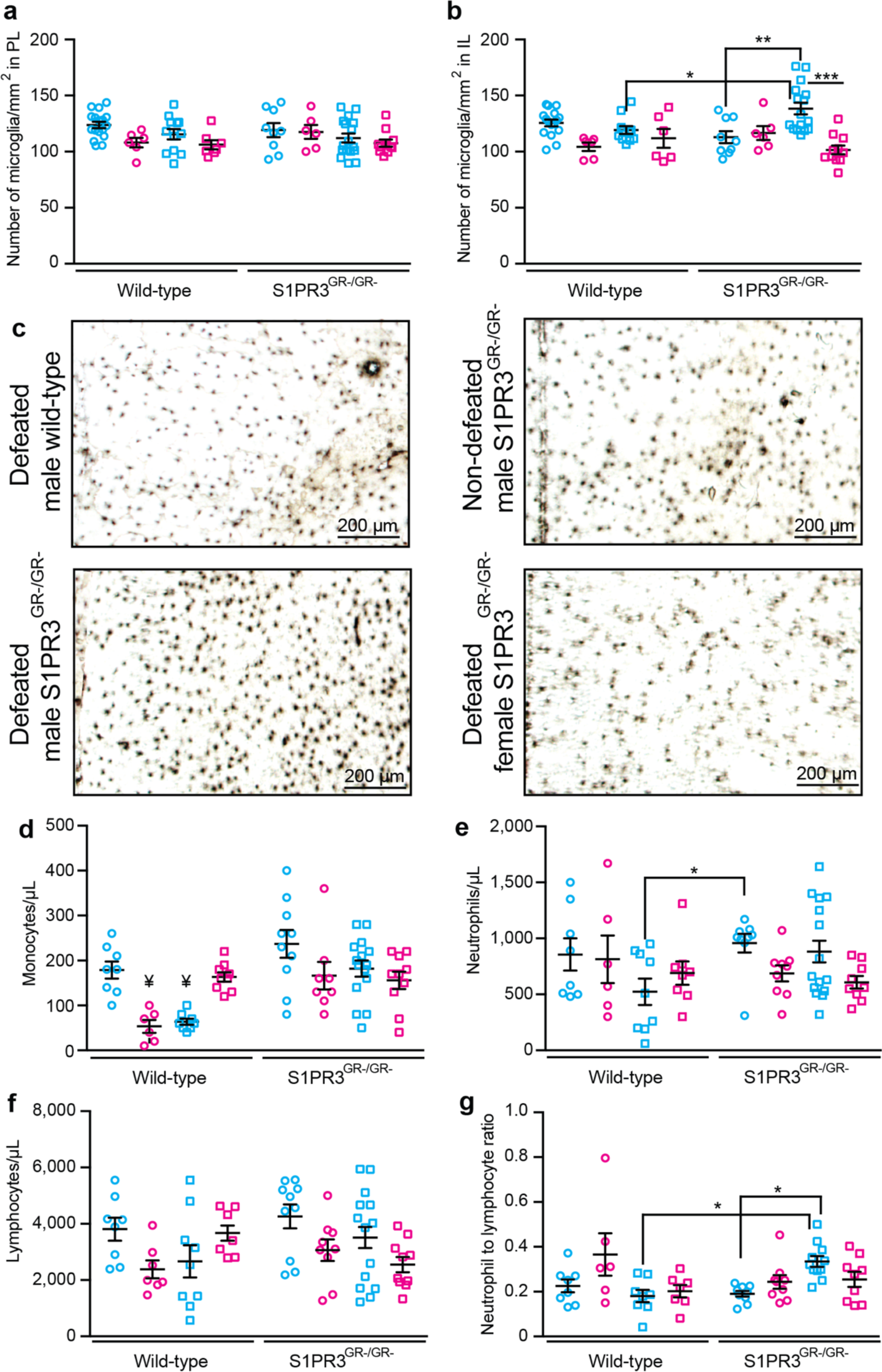
Inflammatory processes are increased in S1PR3^GR-/GR-^ rats. IBA+ cells in the (**a**) PL and (**b**) IL of non-defeated wild-type males (n=16), non-defeated wild-type females (n=6), defeated wild-type males (n=12), defeated wild-type females (n=6), non-defeated S1PR3^GR-/GR-^ males (n=9), non-defeated S1PR3^GR-/GR-^ females (n=6), defeated S1PR3^GR-/GR-^ males (n=16), and defeated S1PR3^GR-/GR-^ females (n=11). (**c**) Images of IBA1+ cells in the IL. (**d**) Monocyte concentrations from non-defeated wild-type males (n=8), non-defeated wild-type females (n=6), defeated wild-type males (n=8), defeated wild-type females (n=9), non-defeated S1PR3^GR-/GR-^ males (n=10), non-defeated S1PR3^GR-/GR-^ females (n=8), defeated S1PR3^GR-/GR-^ males (n=15), and defeated S1PR3^GR-/GR-^ females (n=10). Concentrations of (**e**) neutrophils, (**f**) lymphocytes, and (**g**) the neutrophil-to-lymphocyte ratio in non-defeated wild-type males (n=8), non-defeated wild-type females (n=6), defeated wild-type males (n=9), defeated wild-type females (n=7), non-defeated S1PR3^GR-/GR-^ males (n=9), non-defeated S1PR3^GR-/GR-^ females (n=9), defeated S1PR3^GR-/GR-^ males (n=11), and defeated S1PR3^GR-/GR-^ females (n=9). Lines represent means ± SEM. *p<0.05, **p<0.01, ***p<0.001; Tukey’s multiple comparisons following 3-way ANOVA. For (d), ¥ signifies p<0.05 compared to all groups except one another.

Monocyte concentrations^41^ and neutrophil-to-lymphocyte ratios^42–44^ are higher in humans with stress-related illness, although the underlying mechanisms are not well understood. S1PR3 is expressed in peripheral immune cells and has been shown to reduce inflammatory processes in monocytes^37^. Therefore, we assessed concentrations of circulating monocytes, neutrophils, and lymphocytes in the blood of non-defeated and defeated wild-type and S1PR3^GR-/GR-^ males and females. Monocyte concentrations are lower in non-defeated wild-type females and defeated wild-type males compared to all other groups except defeated S1PR3^GR-/GR-^ females.

This indicates that in females, defeat increases monocyte concentrations and GR-induced S1PR3 maintains low monocyte concentrations under baseline conditions. In males, defeat reduces monocyte concentrations and this requires GR-induced S1PR3 (Fig. 3g). Across all groups, defeat reduced neutrophil concentrations (defeat effect, p = 0.0348) and neutrophil concentrations were higher in non-defeated S1PR3^GR-/GR-^ males compared to defeated wild-type male littermates (Fig. 3h). In general, females displayed lower lymphocyte concentrations compared to males (sex effect, p = 0.0327), but no differences were observed among groups (Fig. 3i). Defeat increased neutrophil-to-lymphocyte ratios in S1PR3^GR-/GR-^, but not wild-type, males and the neutrophil-to-lymphocyte ratio was increased in defeated S1PR3^GR-/GR-^ males compared to defeated wild-type males (Fig. 3j). Together, these findings indicate that GR-induced S1PR3 plays an important role in mitigating stress-induced inflammatory processes in males and in reducing monocyte concentrations in females under baseline conditions.

### mPFC-projecting LC neurons increase LC-mPFC coherence in S1PR3^GR-/GR-^ rats and contribute to stress vulnerability

We previously demonstrated that S1PR3 overexpression in the mPFC reduces c-Fos expression in both the central and basolateral amygdala following defeat^32^, suggesting that S1PR3 stabilizes mPFC efferents during stress. However, it was completely unknown whether S1PR3 is also important for stabilizing stress-induced afferents to the mPFC. Stress increases locus coeruleus (LC) activity and LC-mPFC coherence in the high theta range^33^. To determine whether S1PR3 mitigates the effects of stress-activated mPFC afferents, we assessed local field potentials in the mPFC and LC along with their coherence, a measure of synchronous oscillations, in the delta (1.5-4 Hz), low theta (4-6 Hz), high theta (6-8 Hz), alpha (8-12 Hz), beta (12-20 Hz), and gamma (20-40 Hz) ranges in wild-type and S1PR3^GR-/GR-^ males. To determine whether any changes were driven by mPFC-projecting LC neurons, used Cre-dependent hM4D Designer Receptors Exclusively Activated by Designer Drugs (DREADDs) to chemogenetically inhibit mPFC-projecting LC neurons. 21 days following surgery to express Cre-dependent hM4D and mCherry or mCherry alone, electrodes were implanted in the right LC and mPFC. Rats recovered for seven days and were injected with clozapine-*N*-oxide (CNO), the synthetic ligand for hM4D, 60 minutes prior to defeat onset every day for 7 days. 24 hours later, in the absence of CNO, social interaction was assessed (Figure 4a). The following day rats were euthanized and mCherry expression in the LC was later confirmed (Figure 4b).

**Figure 4.**
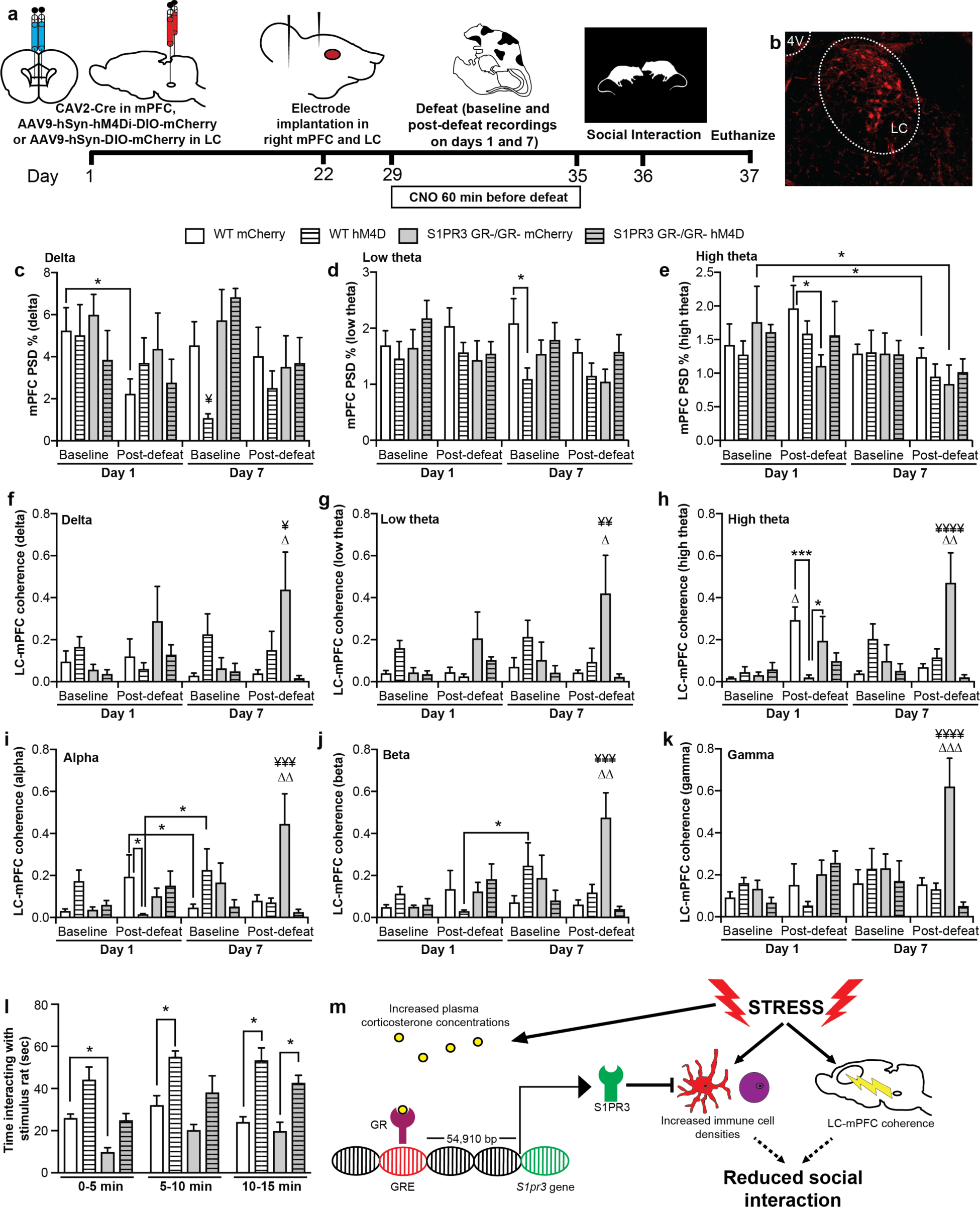
LC-mPFC coherence is increased in defeated male S1PR3^GR-/GR-^ rats and contributes to social anxiety-like behavior. (**a**) Schematic of experimental timeline. (**b**) mCherry expression in mPFC-projecting LC neurons in rats injected with CAV2-Cre in the mPFC and AAV9-hSyn-hM4Di-DIO-mCherry in the LC. Power spectral density percentage in the mPFC in the (**c**) delta, (**d**) low theta, and (**e**) high theta frequency ranges. LC-mPFC coherence in the (**f**) delta, (**g**) low theta, (**h**) high theta, (**i**) alpha, (**j**) beta, and (**k**) gamma frequency ranges. (**l**) Social interaction is reduced in S1PR3^GR-/GR-^ rats (genotype effect p<0.0001) and increased in hM4D-expressing rats (DREADDs effect p<0.0001). (**m**) Graphical summary of results. For all panels and timepoints, wild-type mCherry (n=7), S1PR3^GR-/GR-^ mCherry (n=6), wild-type hM4D (n=6), S1PR3^GR-/GR-^ hM4D (n=5). Lines represent means ± SEM. *p<0.05, **p<0.01, ***p<0.001; ¥p<0.05, ¥¥p<0.01, ¥¥¥p<0.001, ¥¥¥¥p<0.0001 compared to all groups at that timepoint; Δp<0.05, ΔΔp<0.01, ΔΔΔp<0.001 compared to all other timepoints for that group following post-hoc analysis following repeated measures 3-way ANOVA. For (c-k), Fisher’s Least Significant Difference Test was used. For (l), Tukey’s multiple comparisons was used.

In the mPFC, power spectral density percentage (PSD %, power) was reduced by a single defeat in wild-type mCherry rats but not in any other groups. During the day 7 baseline recording, delta power was reduced in wild-type hM4D rats compared to all other groups at that timepoint (Fig. 4c). Compared to mCherry-expressing wild-type controls, hM4D-expressing wild-type rats displayed low theta power in the mPFC (Fig.4d). During post-defeat recordings on day 7, high theta power in the mPFC was reduced in mCherry-expressing S1PR3^GR-/GR-^ rats compared to day 1 baseline recordings and in mCherry-expressing wild-type rats compared to day 1 post-defeat recordings. Following the first defeat, high theta power was reduced in the mPFC of mCherry-expressing, but not hM4D-expressing, S1PR3^GR-/GR-^ rats compared to wild-type controls (Fig. 4e). During the baseline 7 recording, alpha power in the mPFC was increased in hM4D-expessing wild-type rats compared to S1PR3^GR-/GR-^ rats regardless of DREADDs expression (Supplementary Fig. 1c). During the baseline day 7 recoding, beta and gamma power in the mPFC were also increased in hM4D-expressing wild-type rats compared to all other groups at this timepoint (Supplementary Fig. 1d,e). During the post-defeat day 1 recording, low theta power in the LC was decreased in mCherry-expressing S1PR3^GR-/GR-^ rats and trended towards a decrease hM4D-expressing S1PR3^GR-/GR-^ rats compared to their respective wild-type control counterpart. In mCherry-expressing wild-type rats, low theta power in the LC was higher during the day 1 post-defeat recording compared to all other timepoints (Supplementary Fig. 1g). LC power was reduced in the high theta range of mCherry-expressing S1PR3^GR-/GR-^ rats during day 7 baseline and post-defeat recordings compared to their day 1 baseline recording. High theta power and alpha power in the LC was reduced in mCherry-expressing S1PR3^GR-/GR-^ rats compared to mCherry-expressing wild-type controls during the day 7 post-defeat recording (Supplementary Fig. 1h,i). In mCherry-expressing S1PR3^GR-/GR-^ rats, LC alpha power was reduced during the day 7 post-defeat recording compared to the day 1 post-defeat recording (Supplementary Fig. 1i). Together, these findings indicate social defeat, chemogenetic inhibition of mPFC-projecting LC neurons, and GR-induced S1PR3 alter power in specific frequency ranges within the mPFC and LC.

Across all frequency ranges, LC-mPFC coherence was increased during the day 7 post-defeat recording in S1PR3^GR-/GR-^ rats compared to all other groups at that timepoint. In S1PR3^GR-/GR-^ rats, LC-mPFC coherence increased during the day 7 post-defeat recording compared to all other timepoints in all frequency ranges (Fig. 4f-k). LC-mPFC coherence is increased in the high theta range following a single defeat compared to all other timepoints in mCherry-expressing wild-type rats (Fig. 4h). During the day 1 post-defeat recording, LC-mPFC coherence in high theta and alpha was reduced in hM4D-expressing wild-type rats compared to mCherry-expressing wild-type rats (Fig. 4h,i). In mCherry-expressing wild-type rats, LC-mPFC coherence is reduced in the alpha range during the day 7 baseline recording compared to the day 1 post-defeat recording (Fig. 4I). In the alpha and beta frequency ranges, LC-mPFC coherence was increased in hM4D-expressing wild-type rats during the day 7 baseline recording compared to the day 1 post-defeat recording. In sum, these results confirm previous findings that LC-mPFC coherence is increased in the high theta range following a single defeat in wild-type controls^33^. On day 7, when S1PR3 is increased by defeat in wild-type rats, LC-mPFC coherence is increased following defeat in mCherry-expressing S1PR3^GR-/GR-^ rats compared to all other groups. Both of these findings were observed in mCherry-, not hM4D-, expressing rats, indicating that chemogenetic inhibition of mPFC-projecting LC neurons mitigates stress-induced LC-mPFC coherence.

We hypothesized that increased LC input to the mPFC promotes social anxiety-like behavior in wild-type and S1PR3^GR-/GR-^ rats. 24 hours following their 7^th^ defeat, social anxiety-like behavior was assessed in mCherry- and hM4D-expressing wild-type and S1PR3^GR-/GR-^ rats using the social interaction paradigm. Time (p = 0.0013), genotype (p < 0.0001), and DREADDs (p<0.001) effects were observed as social interaction was higher in hM4D-expressing rats regardless of genotype and social interaction was lower in S1PR3^GR-/GR-^ rats regardless of DREADDs. Genotype effects were strongest in the first 5 minutes of the social interaction paradigm whereas DREADDs effects were strongest in 5-10 minute and 10-15 minute intervals (Fig. 4l). This experiment confirms previous findings that social interaction is lower in S1PR3^GR-/GR-^ rats and rats that had S1PR3 knocked down in the mPFC^32^. To the best of our knowledge this finding is the first to demonstrate that chemogenetic inhibition of mPFC-projecting LC neurons during stress increases subsequent social interaction in the absence of any manipulation to neuronal activity. Together, these findings indicate that GR-induced S1PR3 reduces social anxiety-like behavior by mitigating stress-induced inflammatory processes and LC-mPFC coherence (Fig. 4m).

## Discussion

Here we showed that GR-induced S1PR3 reduces inflammatory processes and social anxiety-like behavior in stressed males. These effects were also observed in females under baseline conditions. We hypothesize that effects were observed in females in the absence of stress because baseline corticosterone levels in females are 5-10 higher than those of males^21,23,24^ and similar to plasma corticosterone concentrations observed in stressed males^35^. This caused S1PR3 expression to be higher in the mPFC of wild-type females compared to wild-type males in the absence of stress. This sex difference was not observed in S1PR3^GR-/GR-^ rats, supporting our hypothesis that corticosterone-activated GRs increase S1PR3 in females under baseline conditions. We also showed that LC-mPFC coherence is increased across all frequency ranges during the day 7 post-defeat recording in S1PR3^GR-/GR-^ rats. This indicates that GR-induced S1PR3 mitigates stress-induced increases in LC-mPFC coherence. Indeed, mPFC-projecting LC neurons activated by stress induce social anxiety-like behavior as chemogenetically inhibiting mPFC-projecting LC neurons increased subsequent social interaction in wild-type and S1PR3^GR-/GR-^ rats.

We had previously demonstrated that GRs increase S1PR3 in the mPFC of defeated males^32^. Another group had identified the precise location of the GRE upstream of the gene encoding S1PR3 in mice and reported that no other S1PRs had a proximal GRE^34^. Using the BLAST-Like Alignment Tool (BLAT), we identified a homologous sequence in rats at a comparable distance upstream of the S1PR3 gene. Using ChIP, we confirmed that GRs bind this sequence as GR binding to this locus is increased in defeated wild-type males compared to non-defeated wild-type males. Of course, we did not observe any amplification of this sequence in S1PR3^GR-/GR-^ rats. We did not perform ChIP in females because GR binding is predicted to be similar in non-defeated and defeated females because baseline corticosterone levels are high, resulting in similar expression of S1PR3 in the mPFC of wild-type females regardless of defeat. Indeed, we found that S1PR3 in the PL and IL is higher in non-defeated wild-type females compared to wild-type males. We also found that defeat increases S1PR3 in the IL of defeated males. With the exception of non-defeated males, S1PR3^GR-/GR-^ rats displayed reduced S1PR3 compared to their wild-type counterparts in the PL and IL. In whole blood, we found similar effects under baseline conditions as *S1pr3* mRNA was increased in wild-type females compared to wild-type males and S1PR3^GR-/GR-^ males and females. We also found that *S1pr3* mRNA was reduced in defeated male S1PR3^GR-/GR-^ rats compared to defeated wild-type males. The magnitude of the effect was relatively small, but significant nonetheless. A larger magnitude for this effect might have been observed if *S1pr3* mRNA was assessed in sorted blood immune cells.

Rats that passively cope by displaying shorter defeat latencies exhibit reduced social interaction whereas rats that actively cope by displaying longer defeat latencies behave similarly to non-defeated controls^35^. We previously demonstrated that knocking down S1PR3 in the mPFC reduces defeat latency and overexpressing S1PR3 in the mPFC increases defeat latency in male rats^32^. Indeed, male S1PR3^GR-/GR-^ rats displayed reduced defeat latency compared to wild-type males although this effect was not observed in females. Defeat latencies in females were approximately half those of males, which might be because of a larger size difference between female intruders and residents compared to the size difference in males. It is possible that the effects of GR-induced S1PR3 could not overcome this size difference in females. We previously demonstrated that the effects of S1PR3 on affective-like behavior and inflammatory processes are observed without separating defeated wild-type controls based on defeat latency^32^. We decided to keep defeated rats as one group, rather than segregating them based on defeat latency, because it allowed us to determine overall defeat effects. This would not be possible if we separated defeated rats into short and long latency groups because there cannot be non-defeated counterparts. Although the resident-intruder paradigm of social defeat has been studied in females^45^, the interaction of sex and defeat effects on social interaction and inflammatory processes in the mPFC and blood is not well understood. For these reasons, we felt it was important to avoid separating defeated rats based on defeat latency. Social defeat reduced social interaction in males and females as an overall defeat effect was observed.

Defeated wild-type males, but not females, displayed reduced social interaction compared to their respective non-defeated wild-type controls. The effects of defeat were exacerbated in S1PR3^GR-/GR-^ rats as social interaction was reduced in defeated S1PR3^GR-/GR-^ males and females compared to their defeated wild-type counterparts. Importantly, reduced social interaction in S1PR3^GR-/GR-^ females compared to wild-type females was even observed under baseline conditions, indicating a more severe phenotype in S1PR3^GR-/GR-^ females compared to males. We had previously demonstrated that stress-induced S1PR3 mitigates the development of maladaptive behavior in males. We confirmed this finding here and found that GR-induced S1PR3 may play a more important role in females as it regulates S1PR3-mediated processes, including social anxiety-like behavior and inflammation, even under baseline conditions.

The anti-inflammatory role of S1PR3 in the mPFC^32^ and blood^36,37^, might be attributed to its inhibition of Toll-like receptor 2-mediated nuclear translocation of Nuclear Factor Kappa B (NFkB)^38^. Inflammatory processes might be driven by activation of the coeruleus and sympathetic nervous system as norepinephrine increases microglial reactivity^46^ and cytokine production^47^ in the brain and increases the expression of NFkB in peripheral immune cells^48^. We previously demonstrated that S1PR3 mitigates stress-induced increases in microglia densities and inflammatory cytokines in the mPFC^32^. Consistent with these findings, we report that densities of IBA1+ microglia are increased by defeat in the IL of S1PR3^GR-/GR-^ males. In the PL, a defeat effect was observed and in both the PL and IL significant sex effects are observed.

These findings indicate that IBA1+ cell densities are decreased in the mPFC in females and by defeat, consistent with an anti-inflammatory role of corticosterone^49,50^. Compared to all other groups, monocyte concentrations are reduced in non-defeated wild-type females and defeated males. Because monocyte concentrations were higher in all S1PR3^GR-/GR-^ rats, which indicates that GR-induced S1PR3 mitigates stress-induced increases in monocytes in males and lowers monocyte concentrations in females under baseline conditions. In defeated females, other pro-inflammatory factors caused by stress, such as increased norepinephrine concentrations^48^, might have a greater impact on monocyte proliferation than any S1PR3-mediated reductions. We observed minimal effects on overall neutrophil and lymphocyte concentrations, although significant effects indicated that defeat generally tended to reduce neutrophils and lymphocyte were generally lower in females. We were interested in the neutrophil-to-lymphocyte ratio because it is a biomarker for PTSD^44^ and MDD^42,51^, especially in males^43^. The neutrophil-to-lymphocyte ratio is thought to be a measure of acute inflammatory processes compared to baseline immune function as neutrophils have a half-life of around 6-12 hours^52,53^ whereas the lymphocyte half-life is on the order of weeks^54–56^. We found that the neutrophil-to-lymphocyte ratio was increased in defeated male S1PR3^GR-/GR-^ rats compared to non-defeated S1PR3^GR-/GR-^ males and defeated wild-type males. This suggests that GR-induced S1PR3 mitigates stress-induced neutrophil-to-lymphocyte increases. Although it is not known whether S1PR3 is reduced in MDD patients, S1PR3 mRNA expression is reduced in veterans with post-traumatic stress disorder (PTSD) compared to veterans without PTSD and *S1PR3* mRNA inversely correlates with symptoms of depression (). Together, these findings demonstrate that S1PR3 mitigates stress-induced inflammatory processes in the mPFC and periphery^32^. Further, impaired induction of S1PR3 may at least partially explain increased neutrophil-lymphocyte ratios in certain individuals afflicted with stress-related psychiatric disorders.

S1PR3 couples with Gq and Gi pathways^57–61^, which tend to depolarize and hyperpolarize neurons, respectively^62,63^. S1PR3 increases excitability of hippocampal neurons^64^, although its effects on mPFC output are less clear since S1PR3 overexpression reduces neuronal activity markers in the amygdala, but increases neuronal activity markers in the paraventricular nucleus of the hypothalamus^32^, two brain regions negatively regulated by the mPFC^65,66^. Thus, the effects of S1PR3 on neuronal activity may be cell type- and context-specific. We wanted to understand how GR-induced S1PR3 affected population activity in the mPFC and LC along with their coherence. The LC is the primary source of norepinephrine in the brain^19^, it is activated by social defeat^33^, and it is an important mediator of mPFC function^67,68^.

Previous work demonstrated that LC-mPFC coherence is increased following a single defeat^33^. Coherence is generally interpreted as more effective communication between two brain regions, although coherent oscillations could also be generated by common afferents. We confirmed previous findings that a single defeat increases coherence in the high theta range following a single defeat in wild-type controls. We elaborated on this result by demonstrating that increased LC-mPFC coherence following a single defeat in wild-type controls does not occur when mPFC-projecting LC neurons are chemogenetically inhibited. Therefore, defeat-induced LC-mPFC coherence, at least in the high theta range, can be primarily attributed to mPFC-projecting LC neurons rather than common afferents. Strikingly, LC-mPFC coherence was increased during the day 7 post-defeat recording in mCherry-, but not hM4D-, expressing S1PR3^GR-/GR-^ rats across all frequency ranges. Here, effects were not observed on day 1 because S1PR3 expression is predicted to be similar in wild-type and S1PR3^GR-/GR-^ rats. Effects might only be observed on day 7 because at this timepoint, defeat increases S1PR3 in wild-type, but not S1PR3^GR-/GR-^, rats. Effects might only be observed during the post-defeat recording because LC neurons are activated by stress^33,69^, causing prolonged activation of the mPFC^70,71^. We hypothesize that in wild-type rats, GR-induced S1PR3 mitigates stress-induced LC-mPFC coherence. This increased coherence was not associated with many changes in LFP power in the LC or mPFC during the day 7 post-defeat recording, although LC power in the high theta and alpha ranges was reduced in mCherry-expressing S1PR3^GR-/GR-^ rats compared to mCherry-expressing wild-type controls. Thus, S1PR3 may play a role in desynchronizing oscillations between the LC and mPFC rather than primarily increasing or decreasing their power.

We found that chemogenetically inhibiting mPFC-projecting LC neurons only during social defeat increases subsequent social interaction in wild-type and S1PR3^GR-/GR-^ rats in the absence of any direct manipulations to these neurons during the social interaction paradigm. Chemogenetically inhibiting these neurons during social interaction could have affected other processes, like attention^68^, making it difficult to parse out effects caused by stress. The precise mechanism(s) by which stress-activated mPFC-projecting LC neurons induce social anxiety-like behavior are not known. One potential mechanism is increased inflammatory processes as TNFα is increased in the mPFC of stressed rodents^32,72^ and pharmacological inhibition of TNFα ameliorates anxiety-like behavior^32,73^. Norepinephrine may be an important factor driving stress-induced cytokine production in the mPFC as it increases microglial reactivity and cytokine production in the brain^46,47^. Therefore, increased cytokine expression may represent one important factor caused by stress-induced norepinephrine that causes social anxiety-like behavior. However, other processes mediated by norepinephrine that have continuing effects, such as late phase long-term potentiation^74–76^, may also contribute to lasting effects on social anxiety-like behavior.

We recognize that glucocorticoid signaling regulates neuronal effects that contribute to certain measures of anxiety- and depression-like behavior in rodents^77,78^. These effects include, but are not limited to, promoting dendrite hypertrophy in the amygdala^77^ and dendrite hypotrophy in the hippocampus^79^. However, corticosterone also reduces social anxiety as assessed by the social interaction test in rodents^80^ and increases swimming in the Porsolt forced swim test in females^81^, suggesting an antidepressant-like effect. Therefore, the effects of corticosterone on affective-like behavior are complex and likely depend on the interaction of a variety of factors, including the timeframe and levels of circulating corticosterone, the stress paradigms that are used, the sex of stressed subjects, and the behavioral assays used to measure subsequent behavior. Glucocorticoids and GRs regulate a wide range of functions, including reducing inflammatory processes^82^ that are known to contribute to anxiety- and depression-like behavior in stressed animals^32,73^. We hypothesize that GR-mediated regulation of certain genes, especially those that reduce inflammatory processes like S1PR3, can mitigate certain maladaptive behavioral effects caused by stress.

Future work will focus on the precise epigenetic mechanisms by which GRs regulate S1PR3 expression. Because the GRE is more than 50,000 bp upstream of the *S1pr3* gene, distal interactions involving many proteins are likely required for GR-induced *S1pr3* expression^83^. Among other histone modifications^49,84^, GRs recruit histone acetyl transferases to increase gene expression^85,86^. Other epigenetic mechanisms that increase gene expression, like histone methylation, may also be involved. An important consideration is that deletion of the GRE upstream of the *S1pr3* gene may also modulate the expression of genes other than *S1pr3*. Future work will also investigate the mechanisms by which S1PR3 reduces inflammatory processes and LC-mPFC coherence and the mechanisms by which mPFC-projecting LC neurons reduce subsequent social interaction.

This work is novel for 4 main reasons. First, we show that a specific GRE is important for regulating inflammatory processes and behavior. It is well established that GRs bind to GREs throughout the genome to regulate genes expression^34,84,86^, reduce inflammatory processes^12,49,50^, and modulate brain function and behavior^87–89^, but determining a causative role for GREs can only be accomplished by deleting or mutating them *in vivo.* To the best of our knowledge, this has not been done. CRISPR/Cas9 technology has made this more accessible, which is particularly beneficial in rats since few genetic rat models exist due to the difficulty to altering the rat genome. Second, S1PR3^GR-/GR-^ females display increased circulating monocyte concentrations and reduced social interaction compared to wild-type controls even in the absence of stress. To the best of our knowledge, genetic rat models of stress vulnerability or inflammation that present more severe phenotypes in females have not been reported. Third, we demonstrate that GR-induced S1PR3 mitigates increased LC-mPFC coherence caused by stress. S1PR3 might have important implications for attention and cognitive flexibility, processes that are dependent on the mPFC and LC^67,90^. Fourth, we identify mPFC-projecting LC neurons as a stress-activated projection that contributes to subsequent reductions in social interaction. Together, these findings identify a novel genomic locus that promotes stress resilience by mitigating inflammatory processes and stress-induced coherence in a circuit that contributes to social anxiety-like behavior.

## Acknowledgements

This work was supported by the Defense Advanced Research Projects Agency (DARPA) and the U.S. Army Research Office under grant number W911NF1010093 to SB. Additional support in the form of a Training Grant in Neurodevelopmental Disabilities NIH/NINDS T32 NS007413 and a NARSAD Young Investigator Grant from the Brain & Behavioral Research Foundation (29185) was awarded to BC. The authors have no conflicts of interest.

---

**Supplementary Figure 1.**
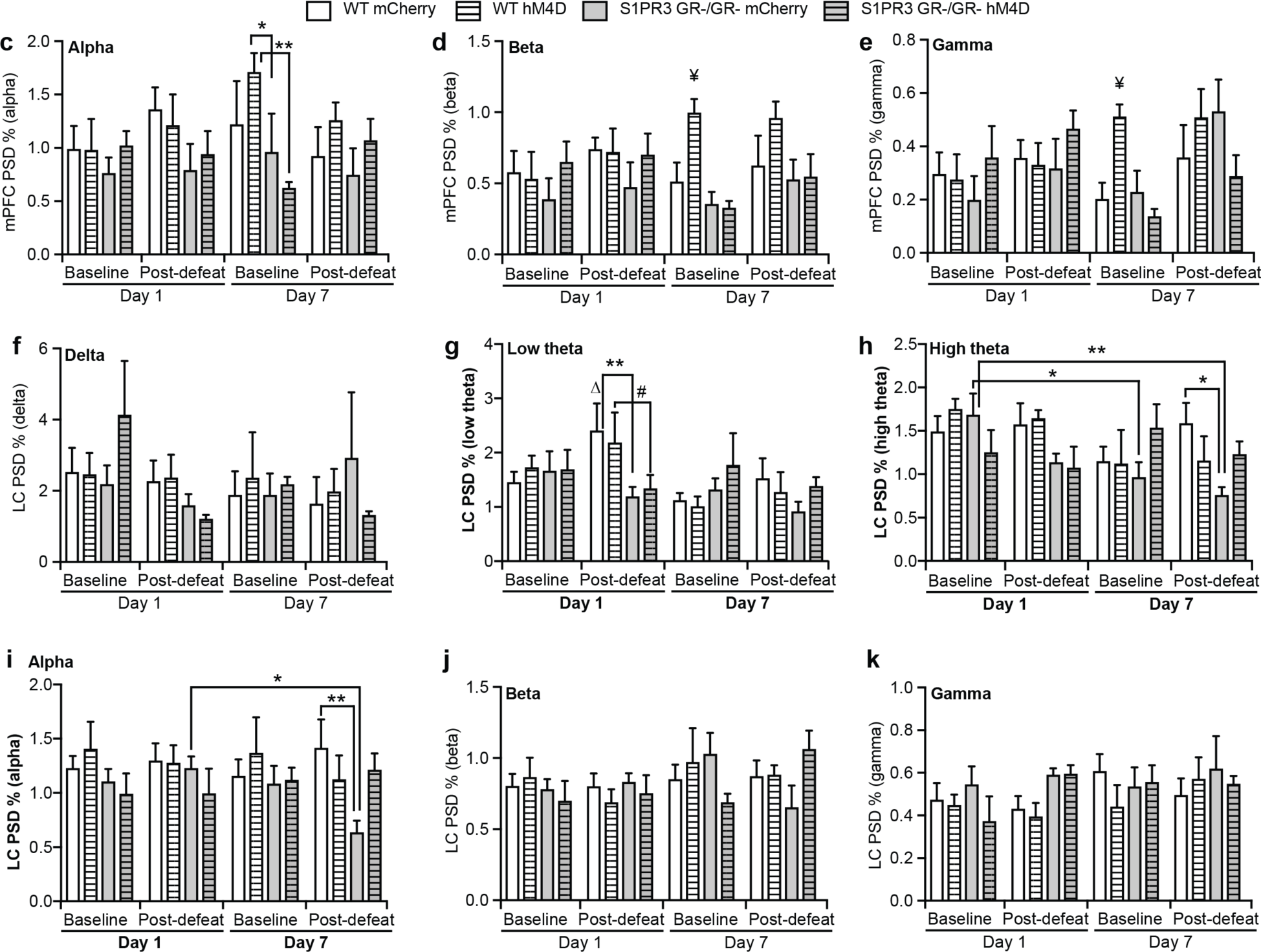
LC-mPFC coherence is increased in defeated male S1PR3^GR-/GR-^ rats and contributes to social anxiety-like behavior. Images of electrode placement in (**a**) the right mPFC (**b**) the right LC. Power spectral density percentages in the mPFC in the (**c**) alpha, (**d**) beta, and (**e**) gamma frequency ranges. Power spectral density percentages in the LC in the (**f**) delta, (**g**) low theta, (**h**) high theta, (**i**) alpha, (**j**) beta, and (**k**) gamma frequency ranges. For all panels and timepoints, wild-type mCherry (n=7), S1PR3^GR-/GR-^ mCherry (n=6), wild-type hM4D (n=6), S1PR3^GR-/GR-^ hM4D (n=5). Lines represent means ± SEM. #p<0.010, *p<0.05, **p<0.01, ¥p<0.05 compared to all groups at that timepoint as assessed by Fisher’s Least Significant Difference Test following repeated measures 3-way ANOVA.

## Online Methods

### Generation of S1PR3^GR-/GR-^ rats

S1PR3^GR-/GR-^ rats were generated under the consultation of the University of Pennsylvania CRISPR/Cas9 Targeting Core. Chromatin immunoprecipitation sequencing had previously identified a GR binding site 34,519 base pairs upstream of the *S1pr3* gene in mice. Notably, 16% of GR binding sites are greater than 50kb from the transcriptional start or stop sites, with only 5% being within 5kb^34^. Using the BLAST-Like Alignment Tool, we identified the homologous sequence in rats. For the generation of S1PR3^GR-/GR-^ rats, a 1,445 base pair sequence on chromosome 17 (5’-13738152-13739596-3’), was deleted. The 3’ end of this sequence is 54,910 base pairs upstream of the first base pair on the 5’ end of the DNA encoding the first exon of the S1PR3 transcript. Guide (g)RNA and Cas9 were microinjected into a fertilized embryo. The 5’ gRNA target sequence was GTGCCCTCTGTAGAACTGCGGGG and the 3’ gRNA target sequence was GAATTGGTCTGTGACGTAGGAGG. The overlapping junction following deletion is AGTGTGCCCTCTGTAGAACTAGGAGGAGCC. Primers used to confirm knockout were CAAGATTCCCAAGAGGATGC (forward) and CCTCAATACAGGGCTCTTGC (reverse). Genotyping was performed by. Four heterozygous founders were created and paired for mating to create homozygous pups. No experiments were performed on any rats until the F4 generations in an effort to avoid any off-target effects.

### Animals

Adult male and female S1PR3^GR-/GR-^ rats and wild-type littermate were used in all experiments. No heterozygotes were used in these studies. Rats were group housed until they were either defeated, served as a novel cage non-defeated control, or underwent surgery. Rats were defeated or served as novel cage controls at 12-14 weeks of age. For experiments requiring virus-mediated gene transfer and *in vivo* electrophysiology, surgery was performed on 8-weel-old rats. Long-Evans retired breeder males were purchased from Charles River Laboratories (Wilmington, MA, USA; 650-850 g) served as residents for male defeat. Residents were singly housed upon arrival. Once experimental procedures began, S1PR3^GR-/GR-^ rats and their wild-type littermates were singly housed in polycarbonate cages with standard bedding and with food and water available *ad libitum*. Animals were kept on a 12-h light–dark cycle with lights on at 06:15 and lights off at 18:15 in a temperature-controlled vivarium for at least 5 days prior to administration of any stress protocols. All experiments took place during the inactive phase between 1000 and 1400 h. Rats were euthanized by rapid decapitation and their brains were immediately snap-frozen in 2-methylbutane. Each day following social defeat, the rats were inspected by the experimenter and an animal technician. Any signs of pain (blood, limping, etc.) were assessed by a veterinarian who recommended euthanasia if symptoms were too severe. Experiments were performed in compliance with all relevant ethical regulations for animal testing and research. Experiment protocols followed the NIH Guide for the Care and Use of Laboratory Animals and were approved by the Children’s Hospital of Philadelphia Research Institute’s Animal Care and Use Committee.

### Chromatin immunoprecipitation

**(ChIP)** ChIP was performed similarly to described methods^21,91–93^ according to the ChIP kit (EMD Millipore, 17-295) manufacturer’s instructions. Prefrontal cortices were subdissected and fixed in 1% formaldehyde. Samples were sonicated to generate genomic fragments 200–1,000 bp in length and pre-cleared with Protein A beads prior to incubation with a rabbit-anti-GR monoclonal primary antibody (Cell Signaling, D6H2L, 12041S) 4°C overnight. The antibody-chromatin complex was immunoprecipitated with Protein A beads, washed with a series of buffers (Millipore), and then chromatin was eluted and reverse cross-linking performed with Proteinase K. DNA was purified via phenol-chloroform extraction. Final DNA concentration was measured using the NanoDrop 2000. qPCR was performed with an ABI 7500 PCR machine using SYBR green as a fluorophore. Primers used to amplify the GRE near the *S1pr3* gene were CTGGAACTGTACCCAACGCT (forward) and AAACTAGCCGAGAAGCCAGG (reverse). The GAPDH primers GACATGCCGCCTGGAGAAAC (forward) and AAGCAGTTGTCCTGTTGGGA (reverse) were used for the housekeeping control.

### Social defeat

The social defeat paradigm was performed as previously described^32,33,35,94^. Rats were randomly assigned to either a social defeat or novel cage control group for 5 to 7 consecutive days. During each episode of social stress, a rat was placed into the home cage territory of an unfamiliar Long-Evans resident previously screened for high aggression. A typical agonistic encounter resulted in intruder subordination or defeat, signaled by the intruder assuming a supine position for 3 sec. After defeat, a wire mesh partition was placed in the cage to prevent physical contact between the resident and intruder but allowing visual, auditory, and olfactory contact for the remainder of the 30 min defeat session. Latency to assume a submissive posture (defeat) was recorded and averaged over the seven daily defeat exposures. Rats that were not attacked were not included in defeat latency analysis for that day. If an intruder resisted defeat for 15 min, the resident and intruder were separated with the wire partition for the remainder of the session. Controls were placed behind a wire partition in a novel cage for 30 min daily. Rats were returned to their home cage after each session.

### Immunohistochemistry

Brains were sectioned on a cryostat at 20 µm. The following primary antibodies were used: rabbit anti-S1PR3/Edg3 (bs-7541R, 1:100, BIOSS), rabbit anti-IBA1 (019-19741, Wako, 1:250), and rabbit anti-mCherry (ab167453, 1:200, Abcam). The following secondary antibodies were used: donkey anti-rabbit (Alexa Fluor ® 488, Abcam, ab150073), donkey anti-rabbit (Alexa Fluor ® 594, Abcam, ab150076), and biotinylated donkey anti-rabbit (Jackson Laboratories, 711-065-152). All secondary antibodies were used at a concentration of 1:200. All immunohistochemical comparisons of protein expression were from assays performed at the same time with the same working solutions. For staining with 3,3’-diaminobenzidine (DAB), further amplification was accomplished using Avidin-Biotin Complex (Vectastain). DAB (Sigma) was used as a chromagen. For quantifying S1PR3, two sections between Bregma + 3.0 and Bregma + 3.4 mm were chosen for analysis. Here, optical density in layer 1 adjacent to the midline was used as a background to be subtracted from neuronal fluorescence. An 8×8 grid was superimposed on each image. S1PR3 immunoreactivity (IR) was quantified in three neurons closest to the grid lines in dorsal, central, and ventral mPFC layers 2/3, 4/5, and 6 (9 cells/section). Means of these cells were created for each section. Sections in which the tissue was not wholly intact or damaged were discarded from analysis.

### Stereotaxic Virus Injections

Cre-dependent AAV9-hSyn-DIO-hM4D-HA-mCherry or control AAV9-hSyn-DIO-HA-mCherry were purchased from the University of North Carolina Vector Core. CAV2-Cre was purchased from *Institut* de *Génétique* Moléculaire de Montpellier, University of Montpellier. Rats were weighed and anesthetized with a ketamine/acepromazine/xylazine cocktail (1/0.2/0.02, 1 mL/kg). The LC (500 nL, A/P: Lamda ™3.7, D/V: 5.1 mm ventral, M/L: ± 1.4 mm) and mPFC (1 µL, A/P: Bregma + 3.2 mm, D/V: 4.4 mm, M/L: ± 0.5 mm) were injected over the course of ten minutes. Surgeries were performed 19-21 days prior to restraint to allow for overexpression. For animal inclusion, mCherry was required to be detected in the LC without significant (>10 cells) expression in other regions in the pons or in the cerebellum. One rat was excluded from the hM4D-expressing S1PR3^GR-/GR-^ group for this reason. Of note, seven S1PR3^GR-/GR-^ rats but only one wild-type control died during surgery due to excessive blood loss. This may be due to a clotting impairment exhibited by S1PR3^GR-/GR-^ rats.

### Social Interaction

Animals were placed in an open field black box (70 cm x 70 cm) with an age-matched stimulus rat of the same strain and of a similar size and allowed to interact for 15 min. Time interacting with the stimulus rat was defined by the time the rat was actively investigating the stimulus rat with its snout closer than 3 cm away (approximately the length of the snout of the rat) from the stimulus rat. Each interaction was videotaped and coded for social interaction time by 2 coders who were blind to the experimental conditions.

### Local field potential recordings

21 days following virus surgery, the right mPFC (A/P: Bregma + 3.2 mm, D/V: 4.4 mm, M/L: 0.5 mm left) and LC (A/P: Lamda ™3.7, D/V: 5.1 mm ventral, M/L:1.4 mm right) had custom recording electrodes (MicroProbes, Part# PI(7mm)0030.1B10) placed within 0.5 mm of the virus injection sites. Cables connected the head stage to the data acquisition system. Rats recovered for 6 days and were subjected to a 10-minute acclimation recording in their home cage. Beginning 24 hours later, rats were injected with clozapine-*N*-oxide (CNO, 2 mg/kg, intraperitoneal) 60 minutes prior to defeat for 7 consecutive days. 10-minute baseline and 10- minute post-defeat recordings were performed on the first and seventh days of defeat. Cables connected the head stage to the data acquisition system. Pre-defeat baseline recordings were done in the intruder’s home cage. The cables were then disconnected and the rat was placed in the resident’s cage for defeat. Post-defeat recordings occurred after the physical interaction, while the intruder was in resident’s cage but was physically separated from the resident by the wire partition, which maintained visual, olfactory, and auditory communication between resident and intruder. Electrode recordings in the mPFC were amplified at a gain of 5000 Hz, bandwidth of 1–150 Hz. Raw mPFC and LC traces were time stamped in Spike2 to remove noise and converted to Power Spectra Density (PSD) plots indicating the relative power and coherence in 128 frequency bins from 0 to 50 Hz using Neuroexplorer (Nex Technologies, Madison, AL).

### Blood collection and isolation of mRNA from whole blood

400 µL of tail blood was collected in RNAprotect Animal Blood Tubes (Qiagen, cat. no. 76554) 24 hours after a 7th day of social defeat. Blood mRNA was isolated using the RNeasy Protect Animal Blood Kit (Qiagen, cat. no. 73224) according to the manufacturer’s instructions.

### qRT-PCR

Reverse transcription was performed using a High-Capacity cDNA Reverse Transcription kit (4368814, Thermo Fischer Scientific). qPCR was performed with an ABI 7500 PCR machine using SYBR Green as a fluorophore. Primers used to amplify cDNA were rat *Gapdh* (forward: 5’-AGACAGCCGCATCTTCTTGT- 3’, reverse: 5’- CTTGCCGTGGGTAGAGTCAT-3’) and rat *S1pr3* (forward: 5’-CCTCATCACCACCATCCTCT- 3’, reverse: 5’- CCCTGAGGAACCACACTGTT-3’) as in our previous work^32^.

### Hematology

90 µL of trunk blood was added to a tube containing 10 µL of ethylenediaminetetraacetic (EDTA). Hematology was performed by the Children’s Hospital of Philadelphia Translational Research Core to quantify concentrations of blood cells using flow cytometry.

### Statistical Analyses

Statistical analyses were performed using Prism 8. Raw means are presented in Supplementary Table 1, significant post-hoc differences are presented in Supplementary Table 2. Differences between means were assessed using a two-tailed, unpaired Student’s t test unless otherwise indicated. Two-way ANOVA with p-corrected Tukey’s multiple comparisons post-hoc testing was used to assess difference among groups in analyses with two factors. Three-way ANOVA with p-corrected Tukey’s multiple comparisons post-hoc testing was used to assess difference among groups in analyses with three factors. Repeated measures was used for analyzing electrophysiological and social interaction data in mCherry- and hM4D-expressing rats. Data beyond three standard deviations from the mean were considered outliers and discarded from analysis.

## References

1 Ehlert, U., Gaab, J., Heinrichs, M. Psychoneuroendocrinological contributions to the etiology of depression, posttraumatic stress disorder, and stress-related bodily disorders: the role of the hypothalamus-piuitary-adrenal axis. Biological Psychiatry 57, 141–152 (2001).

2 Yehuda, R., Teicher, M. H., Trestman, R. L., Levengood, R. A. & Siever, L. J. Cortisol regulation in posttraumatic stress disorder and major depression: a chronobiological analysis. Biol Psychiatry 40, 79–88, doi:10.1016/0006-3223(95)00451-3 (1996).

3 Yehuda, R. Biology of posttraumatic stress disorder. J Clin Psychiatry. 62 Suppl 17, 41–46 (2001).

4 Song, H. et al. Association of Stress-Related Disorders With Subsequent Autoimmune Disease. JAMA 319, 2388–2400, doi:10.1001/jama.2018.7028 (2018).

5 Pace, T. W. & Heim, C. M. A short review on the psychoneuroimmunology of posttraumatic stress disorder: from risk factors to medical comorbidities. Brain, behavior, and immunity 25, 6–13, doi:10.1016/j.bbi.2010.10.003 (2011).

6 O’Donovan, A. et al. Elevated risk for autoimmune disorders in iraq and afghanistan veterans with posttraumatic stress disorder. Biol Psychiatry 77, 365–374, doi:10.1016/j.biopsych.2014.06.015 (2015).

7 Mine, K., Matsumoto, K. & Kanazawa, F. The relation between irritable bowel syndrome and a major depression. Jap. J. Clin. Med. 50, 2719–2723 (1992).

8 Herman, J. Neural control of chronic stress adaptation. Front Behav Neurosci., 61, doi:10.3389/fnbeh.2013.00061. (2013).

9 Sapolsky, R. M., Krey, L.C., & McEwen, B.S. Glucocorticoid-sensitive hippocampal neurons are involved in terminating the adrenocortical stress response. Proc Natl Acad Sci 81, 6174–6177 (1984).

10 Schulman, G. et al. Mineralocorticoid and glucocorticoid receptor steroid binding and localization in colonic cells. Am J Physiol 266, C729–740 (1994).

11 Munck, A., Guyre, P. M. & Holbrook, N. J. Physiological functions of glucocorticoids in stress and their relation to pharmacological actions. Endocrine Rev. 5, 25–55 (1984).

12 Barnes, P. J. & Adcock, I. M. Glucocorticoid resistance in inflammatory diseases. Lancet 373, 1905–1917, doi:10.1016/S0140-6736(09)60326-3 (2009).

13 Dowlati, Y. et al. A meta-analysis of cytokines in major depression. Biol Psychiatry 67, 446–457, doi:10.1016/j.biopsych.2009.09.033 (2010).

14 Rohleder, N., Joksimovic, L., Wolf, J. M. & Kirschbaum, C. Hypocortisolism and increased glucocorticoid sensitivity of pro-Inflammatory cytokine production in Bosnian war refugees with posttraumatic stress disorder. Biol Psychiatry 55, 745–751, doi:10.1016/j.biopsych.2003.11.018 (2004).

15 Yudt, M. R. & Cidlowski, J. A. The glucocorticoid receptor: coding a diversity of proteins and responses through a single gene. Mol Endocrinol 16, 1719–1726, doi:10.1210/me.2002-0106 (2002).

16 Dong, H., Carlton, M. E., Lerner, A. & Epstein, P. M. Effect of cAMP signaling on expression of glucocorticoid receptor, Bim and Bad in glucocorticoid-sensitive and resistant leukemic and multiple myeloma cells. Front Pharmacol 6, 230, doi:10.3389/fphar.2015.00230 (2015).

17 Cattaneo, A. et al. Candidate genes expression profile associated with antidepressants response in the GENDEP study: differentiating between baseline ‘predictors’ and longitudinal ‘targets’. Neuropsychopharmacology 38, 377–385, doi:10.1038/npp.2012.191 (2013).

18 Pariante, C. M. Why are depressed patients inflamed? A reflection on 20 years of research on depression, glucocorticoid resistance and inflammation. Eur Neuropsychopharmacol 27, 554–559, doi:10.1016/j.euroneuro.2017.04.001 (2017).

19 Bangasser, D. A. & Valentino, R. J. Sex differences in stress-related psychiatric disorders: neurobiological perspectives. Front Neuroendocrinol 35, 303–319, doi:10.1016/j.yfrne.2014.03.008 (2014).

20 Rohleder, N., Schommer, N. C., Hellhammer, D. H., Engel, R. & Kirschbaum, C. Sex differences in glucocorticoid sensitivity of proinflammatory cytokine production after psychosocial stress. Psychosom Med 63, 966–972, doi:10.1097/00006842-200111000-00016 (2001).

21 Grafe, L. A., Cornfeld, A., Luz, S., Valentino, R. & Bhatnagar, S. Orexins Mediate Sex Differences in the Stress Response and in Cognitive Flexibility. Biol Psychiatry 81, 683–692, doi:10.1016/j.biopsych.2016.10.013 (2017).

22 Atkinson, H. C. & Waddell, B. J. Circadian variation in basal plasma corticosterone and adrenocorticotropin in the rat: sexual dimorphism and changes across the estrous cycle. Endocrinology 138, 3842–3848, doi:10.1210/endo.138.9.5395 (1997).

23 Figueiredo, H. F., Ulrich-Lai, Y. M., Choi, D. C. & Herman, J. P. Estrogen potentiates adrenocortical responses to stress in female rats. Am J Physiol Endocrinol Metab 292, E1173–1182, doi:10.1152/ajpendo.00102.2006 (2007).

24 Lund, T. D., Munson, D. J., Haldy, M. E. & Handa, R. J. Androgen inhibits, while oestrogen enhances, restraint-induced activation of neuropeptide neurones in the paraventricular nucleus of the hypothalamus. Journal of neuroendocrinology 16, 272–278, doi:10.1111/j.0953-8194.2004.01167.x (2004).

25 Kohler, O. et al. Effect of anti-inflammatory treatment on depression, depressive symptoms, and adverse effects: a systematic review and meta-analysis of randomized clinical trials. JAMA psychiatry 71, 1381–1391, doi:10.1001/jamapsychiatry.2014.1611 (2014).

26 Raison, C. L. et al. A randomized controlled trial of the tumor necrosis factor antagonist infliximab for treatment-resistant depression: the role of baseline inflammatory biomarkers. JAMA psychiatry 70, 31–41, doi:10.1001/2013.jamapsychiatry.4 (2013).

27 Boyle, M. P. et al. Acquired deficit of forebrain glucocorticoid receptor produces depression-like changes in adrenal axis regulation and behavior. Proceedings of the National Academy of Sciences of the United States of America 102, 473–478, doi:10.1073/pnas.0406458102 (2005).

28 Yehuda, R. et al. Cortisol augmentation of a psychological treatment for warfighters with posttraumatic stress disorder: Randomized trial showing improved treatment retention and outcome. Psychoneuroendocrinology 51, 589–597, doi:10.1016/j.psyneuen.2014.08.004 (2015).

29 Suris, A., North, C., Adinoff, B., Powell, C. M. & Greene, R. Effects of exogenous glucocorticoid on combat-related PTSD symptoms. Ann Clin Psychiatry 22, 274–279 (2010).

30 Hori, H. & Kim, Y. Inflammation and post-traumatic stress disorder. Psychiatry Clin Neurosci 73, 143–153, doi:10.1111/pcn.12820 (2019).

31 Yehuda R, Y. R., Buchsbaum MS, Golier JA. Alterations in cortisol negative feedback inhibition as examined using the ACTH response to cortisol administration in PTSD. Psychoneuroendocrinology 31, 447–451 (2006).

32 Corbett, B. F. et al. Sphingosine-1-phosphate receptor 3 in the medial prefrontal cortex promotes stress resilience by reducing inflammatory processes. Nat Commun 10, 3146, doi:10.1038/s41467-019-10904-8 (2019).

33 Zitnik, G. A., Curtis, A. L., Wood, S. K., Arner, J. & Valentino, R. J. Adolescent Social Stress Produces an Enduring Activation of the Rat Locus Coeruleus and Alters its Coherence with the Prefrontal Cortex. Neuropsychopharmacology 41, 1376–1385, doi:10.1038/npp.2015.289 (2016).

34 Yu, C. Y. et al. Genome-wide analysis of glucocorticoid receptor binding regions in adipocytes reveal gene network involved in triglyceride homeostasis. PloS one 5, e15188, doi:10.1371/journal.pone.0015188 (2010).

35 Wood, S. K., Walker, H. E., Valentino, R. J., Bhatnagar, S. Individual differences in reactivity to social stress predict susceptibility and resilience to a depressive phenotype: role of corticotropin-releasing factor. Endocrinology 151, 1795–1805, doi:10.1210/en.2009-1026 (2010).

36 Awojoodu, A. O. et al. Sphingosine 1-phosphate receptor 3 regulates recruitment of anti-inflammatory monocytes to microvessels during implant arteriogenesis. Proceedings of the National Academy of Sciences of the United States of America 110, 13785–13790, doi:10.1073/pnas.1221309110 (2013).

37 Imeri, F. et al. FTY720 and two novel butterfly derivatives exert a general anti-inflammatory potential by reducing immune cell adhesion to endothelial cells through activation of S1P(3) and phosphoinositide 3-kinase. Naunyn Schmiedebergs Arch Pharmacol 388, 1283–1292, doi:10.1007/s00210-015-1159-5 (2015).

38 Dong, Y. F. et al. S1PR3 is essential for phosphorylated fingolimod to protect astrocytes against oxygen-glucose deprivation-induced neuroinflammation via inhibiting TLR2/4- NFkappaB signalling. J Cell Mol Med 22, 3159–3166, doi:10.1111/jcmm.13596 (2018).

39 Barnes, P. J. & Karin, M. Nuclear factor-kappaB: a pivotal transcription factor in chronic inflammatory diseases. The New England journal of medicine 336, 1066–1071, doi:10.1056/NEJM199704103361506 (1997).

40 De Bosscher, K., Vanden Berghe, W. & Haegeman, G. The interplay between the glucocorticoid receptor and nuclear factor-kappaB or activator protein-1: molecular mechanisms for gene repression. Endocr Rev 24, 488–522 (2003).

41 Seidel, A. et al. Major depressive disorder is associated with elevated monocyte counts. Acta Psychiatr Scand 94, 198–204, doi:10.1111/j.1600-0447.1996.tb09849.x (1996).

42 Demir, S. et al. Neutrophil-lymphocyte ratio in patients with major depressive disorder undergoing no pharmacological therapy. Neuropsychiatr Dis Treat 11, 2253–2258, doi:10.2147/NDT.S89470 (2015).

43 Kinoshita, H. et al. Higher neutrophil-lymphocyte ratio is associated with depressive symptoms in Japanese general male population. Scientific reports 12, 9268, doi:10.1038/s41598-022-13562-x (2022).

44 Zhang, M., et al. Relationship between neutrophil to lymphocyte ratio and post-traumatic stress disorder in early stage after acute trauma. Chinese Journal of Emergency Medicine, 479–484 (2021).

45 Reyes, B. A. S. et al. Neurochemically distinct circuitry regulates locus coeruleus activity during female social stress depending on coping style. Brain structure & function, doi:10.1007/s00429-019-01837-5 (2019).

46 Wohleb, E. S. et al. beta-Adrenergic receptor antagonism prevents anxiety-like behavior and microglial reactivity induced by repeated social defeat. The Journal of neuroscience: the official journal of the Society for Neuroscience 31, 6277–6288, doi:10.1523/JNEUROSCI.0450-11.2011 (2011).

47 Blandino, P., Jr., et al. Gene expression changes in the hypothalamus provide evidence for regionally-selective changes in IL-1 and microglial markers after acute stress. Brain, behavior, and immunity 23, 958–968, doi:10.1016/j.bbi.2009.04.013 (2009).

48 Bierhaus, A. et al. A mechanism converting psychosocial stress into mononuclear cell activation. Proceedings of the National Academy of Sciences of the United States of America 100, 1920–1925, doi:10.1073/pnas.0438019100 (2003).

49 Smoak, K. A. & Cidlowski, J. A. Mechanisms of glucocorticoid receptor signaling during inflammation. Mech Ageing Dev 125, 697–706, doi:10.1016/j.mad.2004.06.010 (2004).

50 Barnes, P. J. Corticosteroid effects on cell signalling. Eur Respir J 27, 413–426, doi:10.1183/09031936.06.00125404 (2006).

51 Bustan, Y. et al. Elevated neutrophil to lymphocyte ratio in non-affective psychotic adolescent inpatients: Evidence for early association between inflammation and psychosis. Psychiatry research 262, 149–153, doi:10.1016/j.psychres.2018.02.002 (2018).

52 Summers, C. et al. Neutrophil kinetics in health and disease. Trends Immunol 31, 318–324, doi:10.1016/j.it.2010.05.006 (2010).

53 Rosales, C. Neutrophil: A Cell with Many Roles in Inflammation or Several Cell Types? Front Physiol 9, 113, doi:10.3389/fphys.2018.00113 (2018).

54 Kaur, A. et al. Dynamics of T- and B-lymphocyte turnover in a natural host of simian immunodeficiency virus. J Virol 82, 1084–1093, doi:10.1128/JVI.02197-07 (2008).

55 Ropke, C., Hougen, H. P. & Everett, N. B. Long-lived T and B lymphocytes in the bone marrow and thoracic duct lymph of the mouse. Cell Immunol 15, 82–93, doi:10.1016/0008-8749(75)90166-5 (1975).

56 Hougen, H. P. & Ropke, C. Small lymphocytes in peripheral lymphoid tissues of nude mice. Life-span and distribution. Clin Exp Immunol 22, 528–538 (1975).

57 Windh, R. T. et al. Differential coupling of the sphingosine 1-phosphate receptors Edg-1, Edg-3, and H218/Edg-5 to the G(i), G(q), and G(12) families of heterotrimeric G proteins. J Biol Chem 274, 27351–27358, doi:10.1074/jbc.274.39.27351 (1999).

58 Siehler, S. & Manning, D. R. Pathways of transduction engaged by sphingosine 1- phosphate through G protein-coupled receptors. Biochim Biophys Acta 1582, 94–99, doi:10.1016/s1388-1981(02)00142-7 (2002).

59 Pyne, S. & Pyne, N. Sphingosine 1-phosphate signalling via the endothelial differentiation gene family of G-protein-coupled receptors. Pharmacol Ther 88, 115–131, doi:10.1016/s0163-7258(00)00084-x (2000).

60 Watterson, K., Sankala, H., Milstien, S. & Spiegel, S. Pleiotropic actions of sphingosine-1- phosphate. Prog Lipid Res 42, 344–357, doi:10.1016/s0163-7827(03)00015-8 (2003).

61 Hla, T. Sphingosine 1-phosphate receptors. Prostaglandins Other Lipid Mediat 64, 135–142, doi:10.1016/s0090-6980(01)00109-5 (2001).

62 Roth, B. L. DREADDs for Neuroscientists. Neuron 89, 683–694, doi:10.1016/j.neuron.2016.01.040 (2016).

63 Durkee, C. A. et al. Gi/o protein-coupled receptors inhibit neurons but activate astrocytes and stimulate gliotransmission. Glia 67, 1076–1093, doi:10.1002/glia.23589 (2019).

64 Weth-Malsch, D. et al. Ablation of Sphingosine 1-Phosphate Receptor Subtype 3 Impairs Hippocampal Neuron Excitability In vitro and Spatial Working Memory In vivo. Front Cell Neurosci 10, 258, doi:10.3389/fncel.2016.00258 (2016).

65 Diorio, D., Viau, V., & Meany, M.J. The role of the medial prefrontal cortex (cingulate gyrus) in the regulation of hypothalamic-pituitary-adrenal responses to stress. The Journal of Neuroscience 13, 3839–3847 (1993).

66 Akirav, I. & Maroun, M. The role of the medial prefrontal cortex-amygdala circuit in stress effects on the extinction of fear. Neural plasticity 2007, 30873, doi:10.1155/2007/30873 (2007).

67 Aston-Jones, G., Rajkowski, J. & Cohen, J. Role of locus coeruleus in attention and behavioral flexibility. Biological Psychiatry 46, 1309–1320 (1999).

68 Aston-Jones, G., Rajkowski, J. & Cohen, J. Locus coeruleus and regulation of behavioral flexibility and attention. Prog. Brain Res. 126, 165–182 (2000).

69 Curtis, A. L., Bethea, T. & Valentino, R. J. Sexually dimorphic responses of the brain norepinephrine system to stress and corticotropin-releasing factor. Neuropsychopharmacology 31, 544–554, doi:1300875 [pii] 10.1038/sj.npp.1300875 (2006).

70 Johnson, J. D. et al. Catecholamines mediate stress-induced increases in peripheral and central inflammatory cytokines. Neuroscience 135, 1295–1307, doi:10.1016/j.neuroscience.2005.06.090 (2005).

71 Duran, E., Yang, M., Neves, R., Logothetis, N. K. & Eschenko, O. Modulation of Prefrontal Cortex Slow Oscillations by Phasic Activation of the Locus Coeruleus. Neuroscience 453, 268–279, doi:10.1016/j.neuroscience.2020.11.028 (2021).

72 Audet, M. C., Jacobson-Pick, S., Wann, B. P. & Anisman, H. Social defeat promotes specific cytokine variations within the prefrontal cortex upon subsequent aggressive or endotoxin challenges. Brain, behavior, and immunity 25, 1197–1205, doi:10.1016/j.bbi.2011.03.010 (2011).

73 Karson, A., Demirtas, T., Bayramgurler, D., Balci, F. & Utkan, T. Chronic administration of infliximab (TNF-alpha inhibitor) decreases depression and anxiety-like behaviour in rat model of chronic mild stress. Basic Clin Pharmacol Toxicol 112, 335–340, doi:10.1111/bcpt.12037 (2013).

74 Gelinas, J. N. & Nguyen, P. V. Beta-adrenergic receptor activation facilitates induction of a protein synthesis-dependent late phase of long-term potentiation. The Journal of neuroscience: the official journal of the Society for Neuroscience 25, 3294–3303, doi:10.1523/JNEUROSCI.4175-04.2005 (2005).

75 Straube, T., Korz, V., Balschun, D. & Frey, J. U. Requirement of beta-adrenergic receptor activation and protein synthesis for LTP-reinforcement by novelty in rat dentate gyrus. J Physiol 552, 953–960, doi:10.1113/jphysiol.2003.049452 (2003).

76 O’Dell, T. J., Connor, S. A., Gelinas, J. N. & Nguyen, P. V. Viagra for your synapses: Enhancement of hippocampal long-term potentiation by activation of beta-adrenergic receptors. Cell Signal 22, 728–736, doi:10.1016/j.cellsig.2009.12.004 (2010).

77 Mitra, R. & Sapolsky, R. M. Acute corticosterone treatment is sufficient to induce anxiety and amygdaloid dendritic hypertrophy. Proceedings of the National Academy of Sciences of the United States of America 105, 5573–5578, doi:10.1073/pnas.0705615105 (2008).

78 Gourley, S. L. & Taylor, J. R. Recapitulation and reversal of a persistent depression-like syndrome in rodents. Curr Protoc Neurosci Chapter 9, Unit 9 32, doi:10.1002/0471142301.ns0932s49 (2009).

79 McEwen, B. S. & Sapolsky, R. M. Stress and cognitive function. Current opinion in neurobiology 5, 205–216, doi:10.1016/0959-4388(95)80028-x (1995).

80 File, S. E., Vellucci, S. V. & Wendlandt, S. Corticosterone -- an anxiogenic or an anxiolytic agent? J Pharm Pharmacol 31, 300–305, doi:10.1111/j.2042-7158.1979.tb13505.x (1979).

81 Brotto, L. A., Gorzalka, B. B. & Barr, A. M. Paradoxical effects of chronic corticosterone on forced swim behaviours in aged male and female rats. Eur J Pharmacol 424, 203–209, doi:10.1016/s0014-2999(01)01148-7 (2001).

82 Barnes, P. J. Anti-inflammatory actions of glucocorticoids: molecular mechanisms. Clin Sci (Lond) 94, 557–572, doi:10.1042/cs0940557 (1998).

83 Bartlett, A. A., Lapp, H. E. & Hunter, R. G. Epigenetic Mechanisms of the Glucocorticoid Receptor. Trends Endocrinol Metab 30, 807–818, doi:10.1016/j.tem.2019.07.003 (2019).

84 Ito, K., Barnes, P. J. & Adcock, I. M. Glucocorticoid receptor recruitment of histone deacetylase 2 inhibits interleukin-1beta-induced histone H4 acetylation on lysines 8 and 12. Mol Cell Biol 20, 6891–6903, doi:10.1128/MCB.20.18.6891-6903.2000 (2000).

85 Gaughan, L., Brady, M. E., Cook, S., Neal, D. E. & Robson, C. N. Tip60 is a co-activator specific for class I nuclear hormone receptors. J Biol Chem 276, 46841–46848, doi:10.1074/jbc.M103710200 (2001).

86 Wallberg, A. E., Flinn, E. M., Gustafsson, J. A. & Wright, A. P. Recruitment of chromatin remodelling factors during gene activation via the glucocorticoid receptor N-terminal domain. Biochem Soc Trans 28, 410–414 (2000).

87 McEwen, B. S., Weiss, J. M. & Schwartz, L. S. Selective retention of corticosterone by limbic structures in rat brain. Nature 220, 911–912, doi:10.1038/220911a0 (1968).

88 McEwen, B. S. Plasticity of the hippocampus: adaptation to chronic stress and allostatic load. Ann NY Acad Sci. 933, 265–277 (2001).

89 McEwen, B. S. & Plapinger, L. Association of 3H corticosterone-1,2 with macromolecules extracted from brain cell nuclei. Nature 226, 263–265, doi:10.1038/226263a0 (1970).

90 Park, J. & Moghaddam, B. Impact of anxiety on prefrontal cortex encoding of cognitive flexibility. Neuroscience 345, 193–202, doi:10.1016/j.neuroscience.2016.06.013 (2017).

91 Corbett, B. F. et al. DeltaFosB Regulates Gene Expression and Cognitive Dysfunction in a Mouse Model of Alzheimer’s Disease. Cell reports 20, 344–355, doi:10.1016/j.celrep.2017.06.040 (2017).

92 Renthal, W. et al. Delta FosB mediates epigenetic desensitization of the c-fos gene after chronic amphetamine exposure. The Journal of neuroscience: the official journal of the Society for Neuroscience 28, 7344–7349, doi:10.1523/JNEUROSCI.1043-08.2008 (2008).

93 Tsankova, N. M., Kumar, A. & Nestler, E. J. Histone modifications at gene promoter regions in rat hippocampus after acute and chronic electroconvulsive seizures. The Journal of neuroscience: the official journal of the Society for Neuroscience 24, 5603–5610, doi:10.1523/JNEUROSCI.0589-04.2004 (2004).

94 Wood, S. K., Walker, H. E., Valentino, R. J. & Bhatnagar, S. Individual differences in reactivity to social stress predict susceptibility and resilience to a depressive phenotype: role of corticotropin-releasing factor. Endocrinology 151, 1795–1805, doi:en.2009-1026 [pii] 10.1210/en.2009-1026 (2010).

